# Multiparametric optimization of human primary B-cell cultures using Design of Experiments

**DOI:** 10.1101/2025.01.16.633474

**Authors:** Anne Bruun Rovsing, Kenneth Green, Lisbeth Jensen, Ian Helstrup Nielsen, Jacob Giehm Mikkelsen, Søren E. Degn

## Abstract

B cells are essential in the immune system, driving antibody production, cytokine secretion, and antigen presentation. Studies in mouse models have illuminated key mechanisms underlying B-cell activation, differentiation, class-switch recombination, and somatic hypermutation. However, the extent to which these findings translate to human biology remains unclear. To address this, we developed a human primary B-cell culture system using feeder cells engineered to express CD40L, supplemented with the cytokines BAFF, IL-4, and IL-21. Using a Design of Experiments (DOE) approach, we optimized critical parameters and dissected the individual contributions of each specific factor. Our results reveal that BAFF plays a negligible role, and IL-21 has more subtle effects, whereas CD40L and IL-4 are critical determinants of cell viability, proliferation and IgE class-switching. Furthermore, we find that engineered feeder cells can serve equally well as a source of cytokines, but providing these in purified form increases the flexibility of the system. This platform enables detailed investigation of human B-cell biology, offering insights into intrinsic and extrinsic regulators of antibody responses and providing a foundation for *in vitro* production of human primary antibodies.

## Introduction

B cells are an essential part of our immune defense. They present antigens, produce antibodies, and secrete cytokines. The diversity of the clonally distributed B-cell receptor (BCR) and corresponding secreted antibody repertoire is largely established during B-cell development through somatic gene rearrangement of immunoglobulin (Ig) gene segments encoding the Ig variable regions. Upon activation by antigen, cognate B-cell clones present antigen-derived peptides to CD4+ T-helper cells, proliferate to form a primary focus, and subsequently differentiate through the extrafollicular or germinal center (GC) pathway (Elsner and Shlomchik 2020). Clones may further adapt their BCR through class-switch recombination (CSR), to generate secreted antibody isotypes most relevant for the appropriate immune modular response. Furthermore, in GCs, BCRs can undergo affinity maturation through iterative cycles of clonal expansion with somatic hypermutation (SHM) and competitive selection for increasing antigen affinity. Both CSR and SHM are dependent on the activity of the enzyme activation-induced cytidine deaminase (AID).

*In vitro* cultures attempting to mimic the Thymus (T)-dependent activation of B cells have been performed since the 1990s, where CSR could be induced in purified B cells expressing CD19 using fibroblasts to express CD40 ligand (CD40L) and adding recombinant interleukin-4 (IL-4) (Banchereau and Rousset 1991; Arpin et al. 1995; Fluckiger et al. 1998). In 2009, Luo *et al*. generated antibody-engineered plasma cells by integrating transgenes in hematopoietic stem cells followed by maturation of the cells into antibody-secreting plasma cells (Luo et al. 2009). Following this, Kwakkenbos *et al*. demonstrated that memory B cells can be converted into GC-like B cells by gene transfer of BCL6 and BCL2L1 and co-culture with CD40L-expressing feeder cells, hence enabling continuous culture of antigen-specific B cells (Kwakkenbos et al. 2010). Transgenic BCL6 and BCL2L1 expression resulted in a GC-like state with stable AID expression causing accumulation of Ig heavy-chain variable region (*IGVH)* mutations, resembling SHM. Furthermore, Liu *et al*. developed an *in vitro* culture system enabling the accumulation of mutations resembling SHM using transgenic expression of AID in AID*-*deficient murine B cells stimulated with IL-4 and CD40L (J. Liu et al. 2017). However, the complexity of these systems and the fragile nature of primary B cells have limited the use of these culture systems, and accordingly, primary B-cell cultures have historically not been leveraged to the same extent as primary T-cell cultures (Banchereau 2015).

Using feeder cells expressing CD40L and the recombinant cytokines BAFF, IL-4, and IL-21, Nojima *et al*. developed an *in vitro* culture system capable of driving extensive proliferation, CSR, and differentiation of naïve murine B cells to memory B cells and plasma cells (Nojima et al. 2011). BAFF is an essential survival factor for B cells mainly produced by follicular dendritic cells (FDCs) (Smulski and Eibel 2018), whereas CD40L, IL-4 and IL-21 in combination elicit strong Th2-like T-dependent B-cell activation (Y.-J. Liu et al. 1989; Vercelli et al. 1990; Linterman et al. 2010; Zotos et al. 2010). CD40-ligand (CD40L, gp39, or CD154) is a key co-stimulatory ligand provided by T-helper cells during cognate B-T cell interactions, eliciting a strong activating signal through the NFκB pathway (Elgueta et al. 2009). IL-21 is provided by T-follicular helper (Tfh) cells and is thought to support both proliferation and differentiation of B cells (Ettinger, Kuchen, and Lipsky 2008; Wang et al. 2018). IL-4 is a typical type II immune module cytokine and acts on B cells as an isotype-specifying cytokine towards IgG1 and IgE (Tong and Wesemann 2015).

In further development of the Nojima culture system, Kuraoka *et al*. engineered feeder cells to express BAFF, CD40L, and IL-21. This enabled the expansion of single murine GC B cells into IgG-secreting plasma cells (Kuraoka et al. 2016). Moreover, Su *et al*. developed a system for the differentiation of human naïve B-cells into antigen-presenting B-cells capable of activating T-cells (Su et al. 2016). In recent years, numerous similar culture systems have been established using combinations of these stimuli and additional factors (Caeser et al. 2019; Germar et al. 2019; Koike et al. 2019; Robinson et al. 2019; Finney and Kelsoe 2021; Possamaï et al. 2021; Unger et al. 2021; Marsman et al. 2022; Verstegen et al. 2023). However, the influence of each component and their mutual interaction on B-cell activation, expansion and differentiation remains unclear.

Design of Experiments (DOE) is a structured framework for studying how multiple factors affect a response. DOE can be used to determine whether a list of factors influences the response and whether the factors interact with each other, based on a minimum of experiments. Furthermore, by designing the experiments according to DOE principles, it is possible to build models predicting the response as a function of the factors, which allows for optimization of the response with a minimum of experiments (Tye 2004).

Here, we establish an *in vitro* culture system mimicking the T-dependent activation of human B cells. We leverage the DOE framework to investigate the influence of CD40L, BAFF, IL-4, and IL-21 exposure on human primary B-cell activation, proliferation, and differentiation. This serves as a proof-of-principle for the DOE framework as a viable approach to increase the resolution and transparency of factor contribution for human primary B-cell culture systems and beyond.

## Results

### Generation of a human primary B-cell culture system mimicking T-dependent activation

Inspired by the murine Nojima culture system (Nojima et al. 2011; Kuraoka et al. 2016), we generated feeder cell lines for human CD40L and cytokine delivery by transfection with lentiviral vectors encoding either BAFF, CD40L, IL-4 or IL-21 (**Fig. 1A, B**). We chose primary Normal Human Dermal Fibroblasts (NHDFs) and a murine stromal cell line (MS-5) as feeder cells as they show contact inhibition, hence avoiding the need for irradiation, and furthermore, we had previously seen exceptional slow growth rates for two donors of NHDFs, NHDF03 and NHDF15 (Haslund et al. 2019). The primary NHDFs were immortalized by lentiviral gene transfer of human Telomerase Reverse Transcriptase (hTERT) and were found to sustain culture for at least 4 months without showing any signs of growth arrest. NHDF15 had the slowest doubling time at 64 hours compared to NHDF03 at 48 hours (**Fig. 1C, D**).

**Figure 1.**
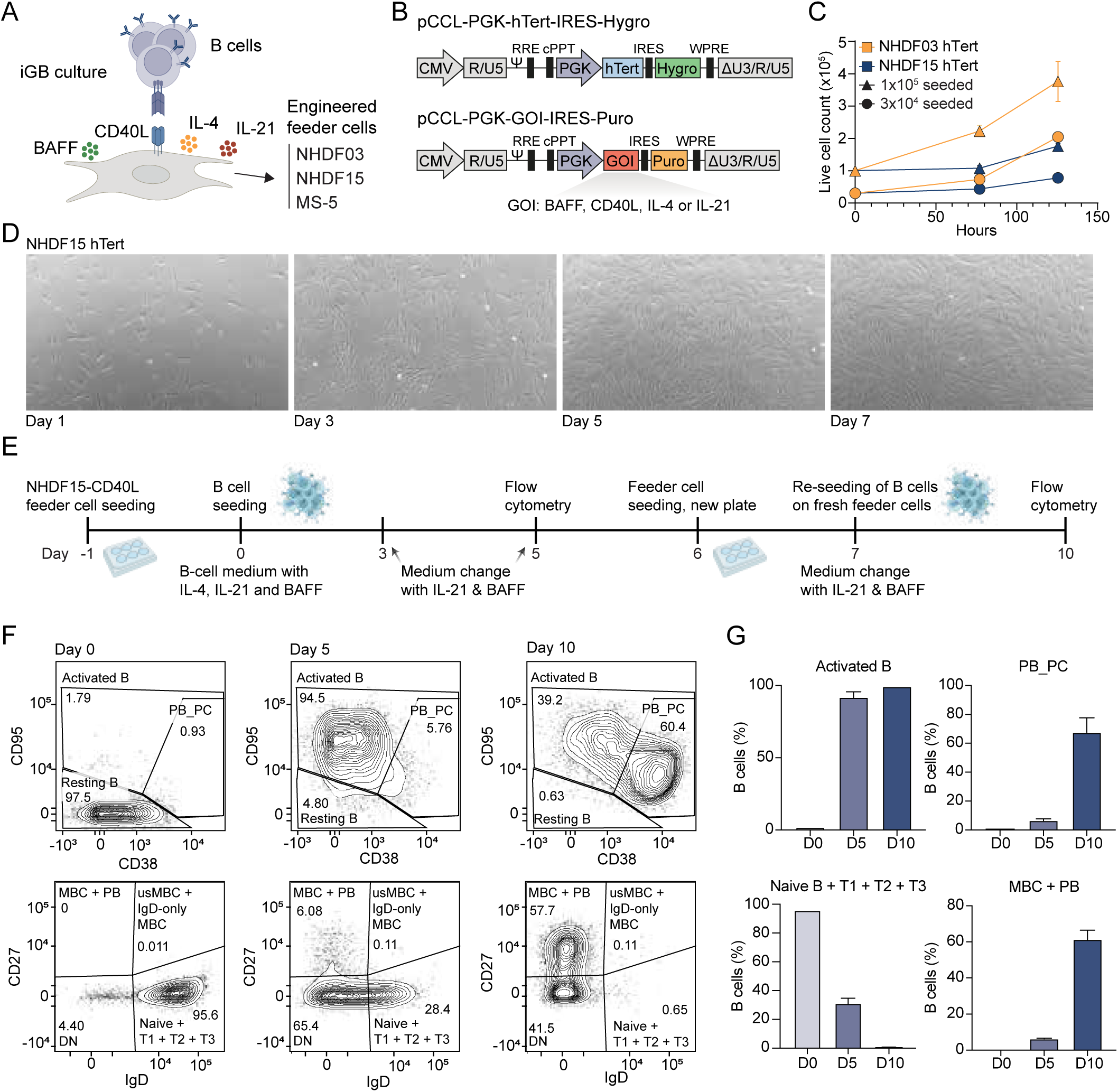
Establishment of a human primary B-cell co-culture system. (**A**) Schematic of the iGB co-culture. Human naïve B cells are seeded on feeder cells expressing CD40L with the cytokines BAFF, IL-4 and IL-21 to promote B-cell activation and differentiation. (**B**) Lentiviral vectors used for engineering feeder cells with human telomerase (hTert), CD40L, BAFF, IL-4, and IL-21. (**C**) Growth curves for hTert-engineered NHDF03 and NHDF15 fibroblasts. Cells were seeded at 1×10^5^ (triangles) or 3×10^4^ (squares) per well in 6-well plates. (**D**) Phase-contrast microscopy images (10x) showing the morphology of NHDF15-hTert cells at different time points. (**E**) Timeline of iGB culture performed in 6-well plates with harvest of cells on day 0, 5 and 10 for flow cytometry. NHDF15-hTert-CD40L feeder cells (1×10^5^) were seeded on day -1 and day 6. Naïve B cells (2.8×10^4^) were seeded on day 0 and re-seeded on fresh feeder cells at day 7. (**F**) Flow cytometric phenotyping of naïve B cell activation and differentiation over time in iGB cultures. (**G**) Frequency of subpopulations from flow cytometry (n=2). Error bars denote mean ± S.E.M.

To test the B-cell activation capability of the engineered feeder cells, human primary naïve B cells were seeded onto the CD40L-expressing feeder cell layer in BAFF, IL-4 and IL-21 cytokine-supplemented medium (**Fig. 1E**). Analyzing by flow cytometry, all three CD40L-expressing feeder cell lines were capable of supporting naïve B cells to become activated B cells (CD38^lo^CD95^hi^) on day 5 and memory-like and plasmablast-like B cells (IgD^−^CD27^+^) on day 10 (data shown for NHDF15-CD40L in **Fig. 1F-G, Fig. S1**; for NHDF03 in **Fig. S2**; data for MS-5-CD40L can be found in (Nielsen et al. 2024).

We generated clones for each of the three feeder cell types (NHDF03, NHDF15, and MS-5) with each of the four transgenes and measured the membrane display of CD40L (**Fig. S3A**) and secretion of BAFF, IL-4, and IL-21 (**Fig. S3B-D**). For NHDF15, we chose the clones with the highest cytokine secretion levels as well as high and low CD40L-expressing clones and further characterized these clones in terms of their proliferation and stability. After multiple freeze-thaw cycles, the NHDF15-CD40L low-expressing clone 2 (N.40-low) and high-expressing clone 10 (N.40-high) showed stable expression levels by flow cytometry (**Fig. S4A**). Likewise, we tested the stability of NHDF15-BAFF clone 1 (N.BAFF), NHDF15-IL-4 clone 3 (N.IL4), and NHDF15-IL-21 clone 1 (N.IL21) and found these to be stable across 30 days of continuous culture by measuring their transgenic cytokine secretion levels by ELISA (**Fig. S4B**). Furthermore, we measured the proliferation rate of each clone and with the exception of N.BAFF each clonal cell line had a proliferation rate comparable to or slower than the parental NHDF15-hTert cells (**Fig. S4C**), and all generated cell lines continued to show contact inhibition.

In summary, we successfully generated CD40L-expressing and cytokine-secreting cell lines with stable expression and slow growth rates, making them suitable for co-cultures without the use of irradiation or cytostatic agents. Furthermore, we demonstrated that the CD40L-expressing cells could support the activation and differentiation of human primary B cells.

### Using DOE to characterize the influence of BAFF, CD40L, IL-4, and IL-21 on B-cell cultures

To investigate the influence of each of the components, BAFF, CD40L, IL-4, and IL-21, we used the DOE framework to enable mathematical modelling of the effect of each component. In the first DOE, we seeded N.40-low cells in 96-well plates on day –1, followed by CD19+ B cells on day 0, and then measured the viability and proliferation of the cultured B cells on day 7 using flow cytometry (**Fig. 2A**). We tested different concentrations of recombinant BAFF, IL-4, and IL-21 as well as different densities of N.40-low (**Fig. 2B**). This was tested as a Full Factorial DOE design entailing two levels (- 1 and 1) of each component being tested for every possible combination, as shown for a three-component mock example (**Fig. 2C**). We then created a linear model with a variable for every component and every interaction between each component and performed Backward Elimination such that only variables with a statistically significant (p < 0.05) coefficient were kept in the model. All DOE experiments and their linear models can be found as R scripts on GitHub (https://github.com/arovsing/DOE_iGB-culture).

**Figure 2.**
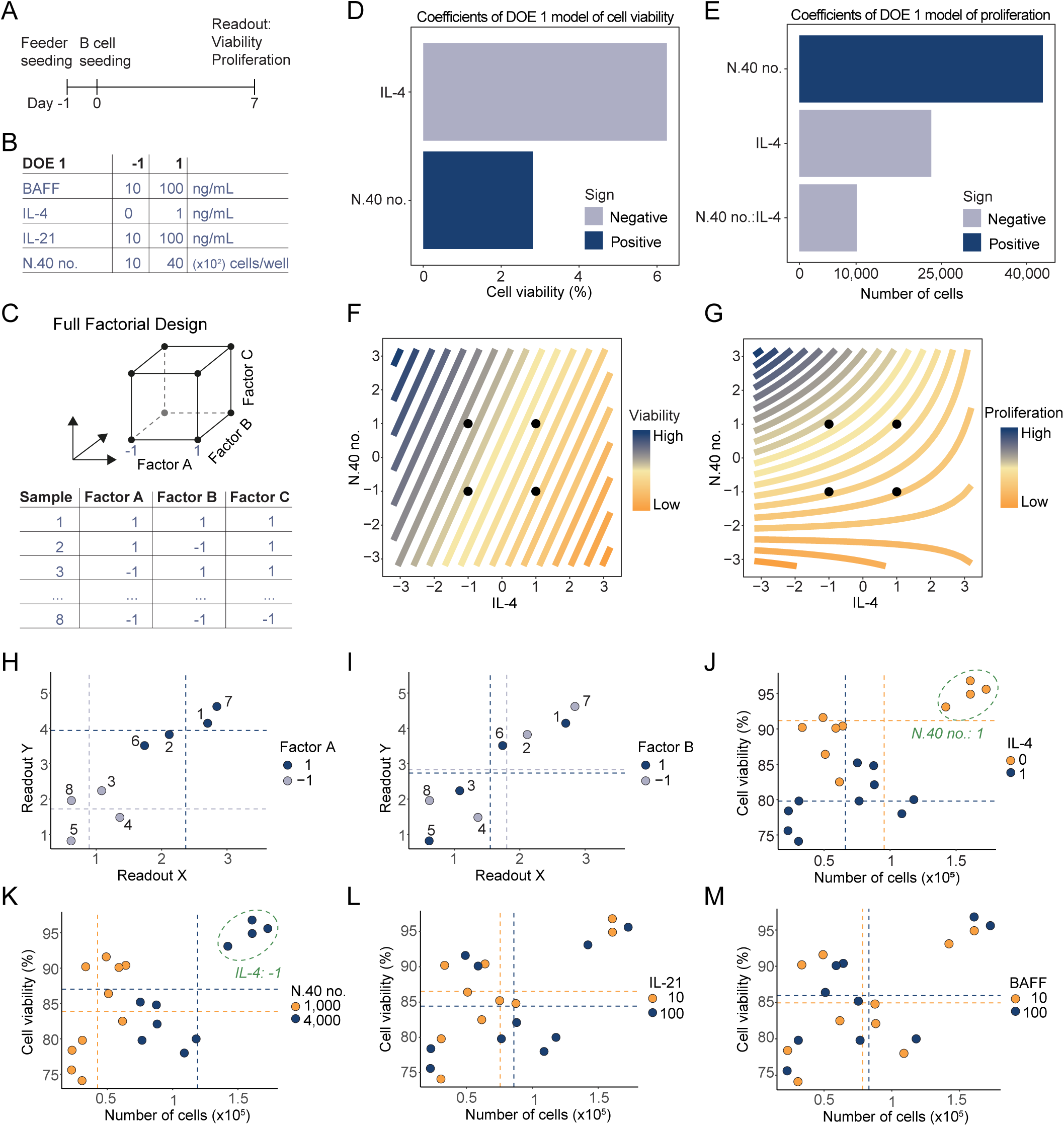
IL-4 and CD40L are critical for B-cell viability and proliferation in iGB cultures. (**A**) Timeline of DOE 1. B-cells (1×10^3^) were seeded in 96-well plates on day 0. Flow cytometry was performed on day 7. (**B**) Design table for DOE 1 showing the high (+1) and low (−1) levels of each tested factor. (**C**) Schematic of a three-factor full factorial DOE design. Each experiment corresponds to a corner of the cube. (**D-E**) Pareto plots of the effect size of different factors on B-cell viability (**D**) and proliferation (**E**) at day 7 of iGB culturing. (**F-G**) Contour plots showing the effect of IL-4 and number of N.40 feeder cells on viability (**F**) and proliferation (**G**). Curved lines indicate an interaction between the two factors. (**H-I**) Conceptual illustration of plotting the data for a three-factor mock example. Each dot represents a sample, and the dotted line represents the mean of all samples with the color-indicated factor value. (**J-M**) Proliferation and viability of each sample from DOE 1 where color and mean indicated by dotted lines are specific for IL-4 (**J**), number of N.40 feeder cells (**K**), IL-21 (**L**), and BAFF (**M**).

For DOE 1, the statistically significant variables at the concentrations tested were IL-4 and the seeding density of N.40-low, of which a high concentration of IL-4 had a negative impact on viability, whereas a high cell density of N.40-low had a positive impact on proliferation (**Fig. 2D, E**). Furthermore, the model for B-cell proliferation showed a negative interaction between the addition of IL-4 and the cell density of N.40-low suggesting that there is a negative synergistic effect between a high concentration of IL-4 together with a high cell density of N.40-low (**Fig. 2F, G**). As interpreting models with many interaction terms is not intuitive, we also plotted the raw data color-coded to each component’s level to provide a visual overview of each component’s effect. In these plots, we display each sample and the level of the specific components, *e.g.* for Factor A and Factor B in the mock example (**Fig. 2H, I**). We also display the mean for each readout for each level, making it possible to see on the raw data if the different levels of a component significantly change the readout. For IL-4 in DOE 1, it was evident that the addition of 1 ng/mL IL-4 led to a significant drop in the viability of more than 10% (**Fig. 2J**), whereas, for N.40-low, a cell density of 4,000 resulted in more than twice as many B cells compared to an N.40-low density of 1,000 (**Fig. 2K**). It was also apparent that at the tested concentrations (10-100 ng/mL) of BAFF and IL-21, neither of these significantly impacted the readout responses (**Fig. 2L, M**).

Altogether, by using the DOE framework we observed that IL-4 negatively impacted viability and CD40L supported proliferation, and at the tested concentrations (10-100 ng/mL) of BAFF and IL-21, these factors did not significantly impact B-cell viability or proliferation.

### DOE characterization of culture system where feeder cells deliver the cytokines

Next, we wanted to test if by delivering the cytokines using feeder cells, we could achieve the same degree of B-cell activation. For DOE 2, we tested different seeding densities of N.BAFF, N.IL4, N.IL21, N.40, and either using N.40-low or N.40-high (**Fig. 3A**). In the linear models for both viability and proliferation, IL-4 was the most significant variable with a negative effect on both, followed by the CD40L expression level and seeding density of N.40 (**Fig. 3B, C**). Modelling both the number of N.40 cells seeded and the CD40L expression level showed opposite effects between the two factors for proliferation. The feeder cell seeding density had a positive effect, whereas the CD40L expression level had a negative effect, and the interaction between the two was negative. This suggests that it is more beneficial for the B-cell proliferation to have many feeder cells with low CD40L expression than having fewer feeder cells with high CD40L expression. Interestingly, upon phase-contrast microscopy, we observed a difference in the appearance of the activated B cells, which displayed a higher tendency to appear in small clusters for N.40-high compared to N.40-low (**Fig. 3D**).

**Figure 3.**
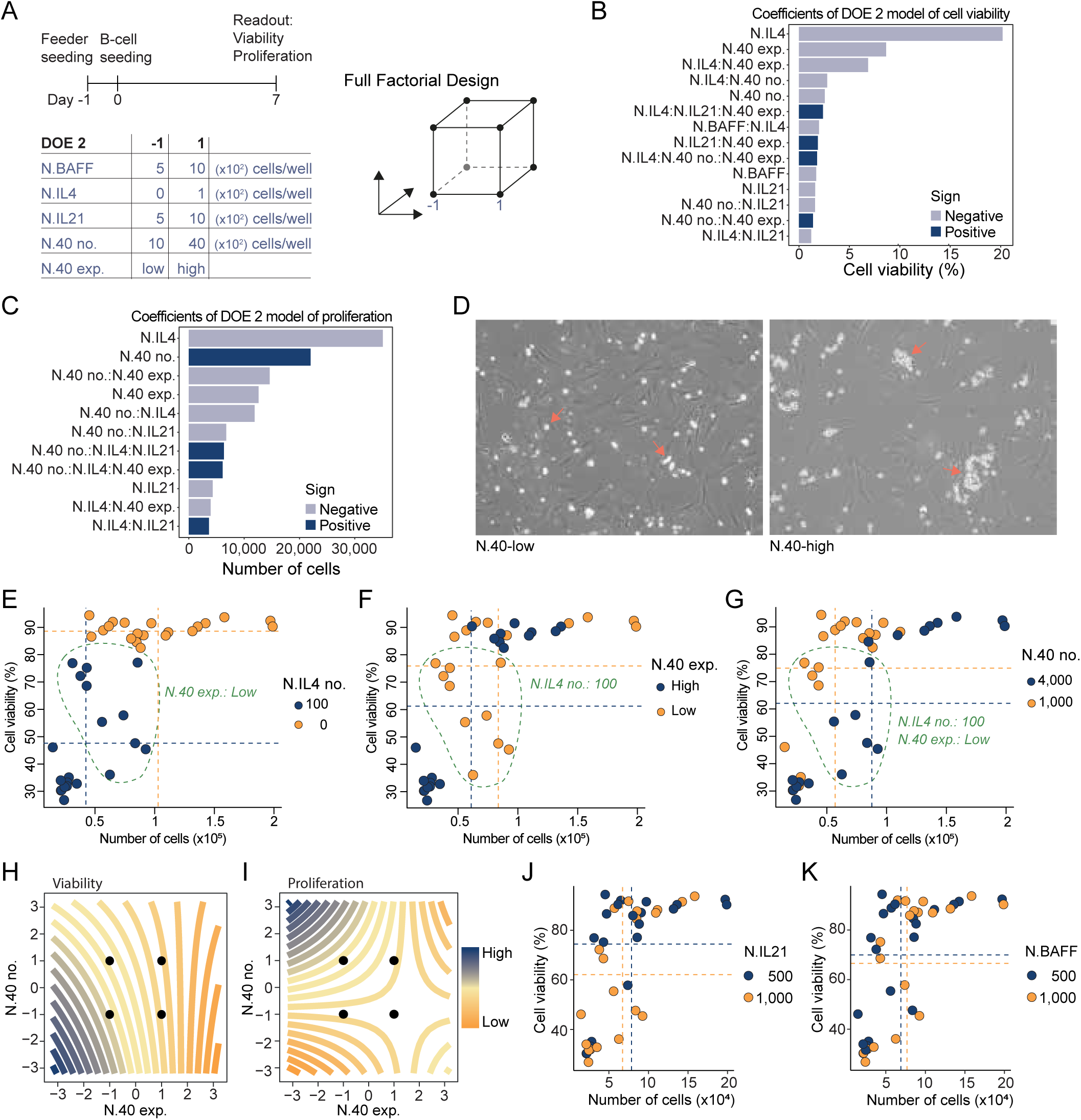
Feeder-cell delivery of cytokines promotes a similar level of B-cell activation as soluble factors. (**A**) Timeline and design table for DOE 2. B-cells (1×10^3^) were seeded in 96-well plates on day 0. Flow cytometry was performed on day 7. (**B-C**) Pareto plot of factor effect sizes on B-cell viability (**B**) and proliferation (**C**). (**D**) Representative phase-contrast microscopy images (10x) of iGB cultures using N.40-low (left) vs. N.40-high (right) feeder cells. N.40 high show more clustering of B-cells. (**E-G**) Proliferation and viability of each sample from DOE 2 where color and mean indicated by dotted lines are specific for number of N.IL4 cells (**E**), level of expression of N.40 cells (**F**), number of N.40 cells (**G**). (**H-I**) Contour plot showing effect on B-cell viability (**H**) and proliferation (**I**) of CD40L expression level of feeder cells compared to their respective seeding density. (**J-K**) Proliferation and viability of each sample from DOE 2 where color and mean indicated by dotted lines are specific for number of N.IL21 cells (**J**) and N.BAFF cells (**K**).

The raw data color-coded to each factor exposed the drastic effect of N.IL4 where the seeding of as few as 100 N.IL4 cells compared to no N.IL4 reduced the viability and proliferation on average by more than 40% and 50%, respectively (**Fig. 3E**). Furthermore, for the B cells cultured with N.IL4, the expression level of N.40 divided the samples into two distinct clusters with the N.40-low resulting in both higher viability and proliferation than N.40-high (**Fig. 3F**). Moreover, for the samples seeded with N.IL4 and N.40-low, the seeding density of N.40 further divided the samples into two distinct clusters with the low seeding density resulting in higher viability, but lower proliferation of B cells indicating the interaction effects between these 3 factors (**Fig. 3G**). On the contrary, for the B cells cultured without N.IL4, neither the CD40L expression level nor N.40 density affected the viability.

Contour plots based on the full models showed that for viability, the optimal condition was to have both low feeder cell density and low CD40L expression, whereas for proliferation it was best to have high density and low CD40L expression (**Fig. 3H, I**). In addition, as we had seen in the setup with purified recombinant cytokines, the effects of N.BAFF and N.IL21 were insignificant (**Fig. 3J, K**).

In summary, we observed that for both B-cell proliferation and viability it was best not to include N.IL4 and use the CD40L low-expressing N.40 feeder cells. The density of N.40 showed different effects with a low density being best for viability and a high density being best for proliferation. As previously seen in the setup with recombinant cytokines, neither N.BAFF nor N.IL21 had a significant effect at the tested densities (5-10 ×10^3^ cells/well).

### Expanding the DOE design to include measurements of Ig-secretion

To further investigate the B-cell culture system, we incorporated a harvest of supernatants from the cultures and performed solid-phase assays (TRIFMA) for IgG and IgE secretion. Moreover, we investigated a broader range of concentrations using a Box Behnken design with 3 levels and non-linear model terms to enable estimation of the optimum concentration. Lastly, to further investigate the influence of IL-4, we incorporated a change of half the medium on day 5 after B-cell seeding with no replenishment of IL-4 as a component in our DOE design (**Fig. 4A**).

**Figure 4.**
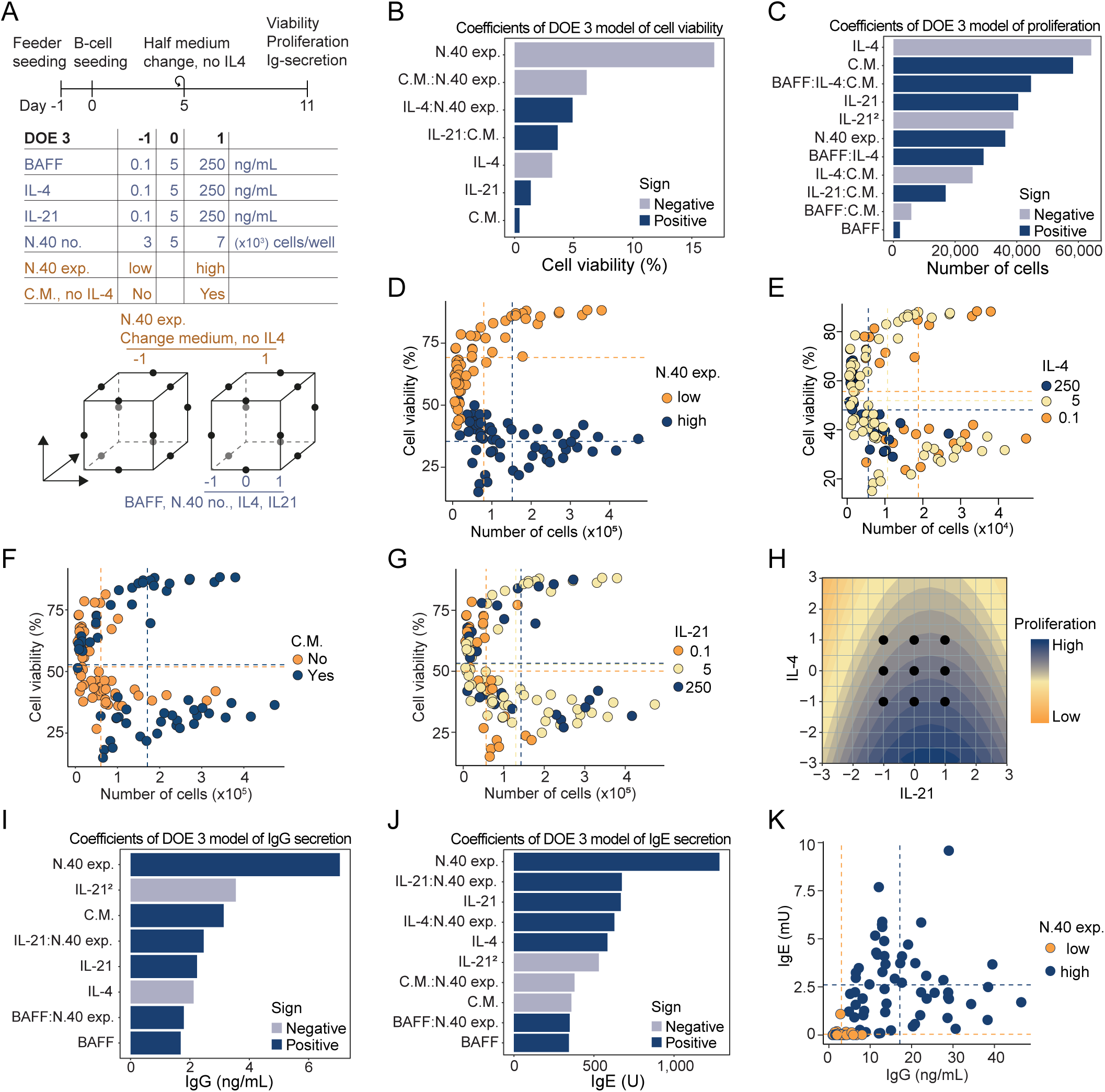
Optimization of iGB cultures using a Box-Behnken DOE strategy. (**A**) Timeline and design table for DOE 3 using a three-level Box-Behnken design. B-cells (1×10^3^) were seeded in 96-well plates on day 0. Flow cytometry and TRIFMA were performed on day 11. (**B-C**) Pareto plots of the effect size of each factor on B-cell viability (**B**) and proliferation (**C**) after 11 days of iGB culturing. IL21^2^ indicates a non-linear contribution from IL-21. (**D-G**) Proliferation and viability of each sample from DOE 3 where color and mean indicated by dotted lines are specific for level of expression of N.40 cells (**D**), IL-4 (**E**), change of medium (**F**), and IL-21 (**G**). (**H**) Contour plot showing effect of IL-21 and IL-4 exposure levels on B-cell proliferation. (**I-J**) Pareto plots of parameter effect sizes on the IgG (**I**) and IgE (**J**) levels measured in culture medium using TRIFMA. (**K**) Measured IgG and IgE in culture medium, color-coded for the use of high (N.40-high) or low (N.40-low) CD40L-expressing feeder cells. C.M. indicates medium change.

As we had seen previously, the factors CD40L expression and IL-4 concentration were the most influential regarding B-cell viability and proliferation (**Fig. 4B-E**). The third most important component was change of half the medium with no replenishment of IL-4 on day 5 (**Fig. 4F**). The level of BAFF and N.40 density in the range tested here (3-7 ×10^3^ cells/well) both had a negligible influence (**Fig. S5A-B**), although BAFF did show signs of interaction with IL-4 and the change of medium in the model for proliferation (**Fig. 4C**). In terms of finding the optimal concentration, the optimal IL-4 concentration was at the lowest level, *i.e.*, 0.1 ng/mL. However, for IL-21 the model for proliferation suggested that the optimal concentration of IL-21 was at level 0.7 translating to 172 ng/mL (**Fig. 4G-H**).

Modelling the IgG and IgE secretion showed that the expression level of CD40L was by far the most influential factor with N.40-high leading to 4-fold and 10-fold higher levels of IgG and IgE detected in the supernatant at day 11 (**Fig. 4I-K**). Besides this, the concentration of IL-4, IL-21 and the change of medium with no IL-4 replenishment also had significant effects on the secretion levels with higher levels of IL-21 leading to more secretion (**Fig. S5C-E**). For IL-4 and change of medium there was an interesting trend with high levels of IL-4 and no removal of IL-4 by medium exchange, where presence of IL-4 led to more IgE secretion, whereas the opposite was true for secretion of IgG. Notably neither BAFF nor the applied seeding-density range (3-7 ×10^3^ cells/well) of N.40 showed any substantial influence on Ig-secretion (**Fig. S5F-G**).

Taken together, these findings confirmed the effect of IL-4 and CD40L on viability and proliferation with high IL-4 being detrimental and low CD40L being beneficial. Conversely, a high IL-4 level supported IgE class-switching while limiting IgG, and a high CD40L level supported class-switching in general. Finally, we demonstrated that the DOE approach could determine optimal concentrations of IL-4 and IL-21 using a minimized experimental setup.

### DOE setup with feeder cell secretion comparable to cytokine delivery

To further investigate if we could achieve the same degree of B-cell activation during culture with feeder cells secreting the cytokines, we set up another round of DOE experiments, in which we varied cytokine concentrations (DOE4) or feeder cell seeding densities (DOE5) (**Fig. 5A**). Here, we changed the IL-4 and BAFF components to be either included or excluded. As before, the addition of IL-4 was by far the most significant component with a markedly higher cell viability and proliferation seen for the samples without IL-4 (**Fig. 5B-C and Fig. S6A**). As previously seen for the samples receiving IL-4, a low CD40L expression level of the feeder cells resulted in a significantly higher viability and proliferation rate of the B cells (**Fig. 5B-D and Fig. S6A**). In DOE 5, where BAFF and IL-21 were secreted by feeder cells, we observed the same trend with IL-4 being the most significant component, and with an interaction between the CD40L expression level and IL-4 (**Fig. 5E-G and Fig. S6B**). For the secretion of IgG and IgE, we similarly observed that IL-4 and the expression level of CD40L were the most impactful components both for DOE 4 and DOE 5 (**Fig. 5H-M and Fig. S6C-D**). In fact, IgE was only secreted if IL-4 was added to the culture system with the lower expressing N.40-low cell line (**Fig. 5I-J, L-M**).

**Figure 5.**
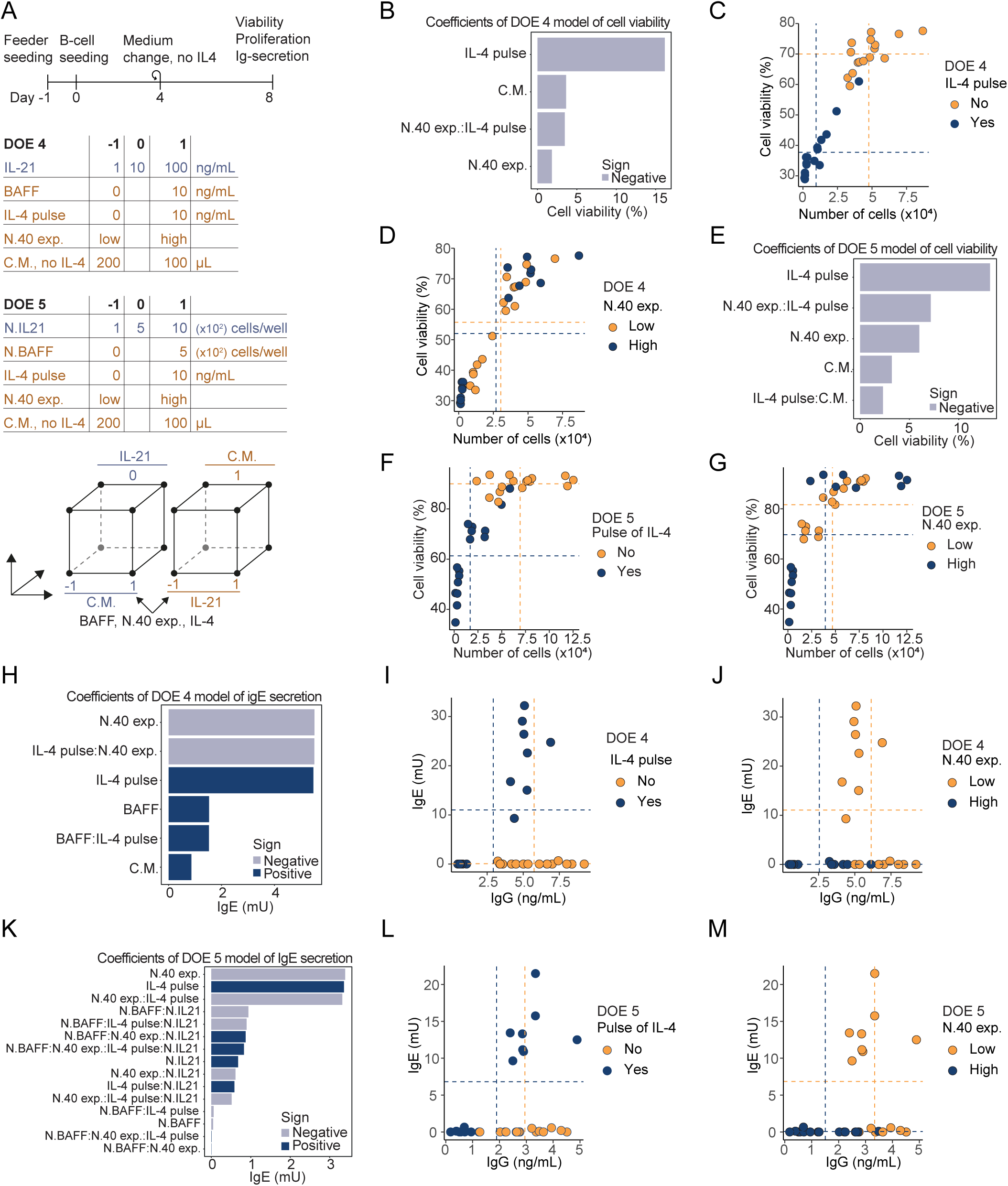
Comparing cytokine delivery methods for iGB cultures. (**A**) Timeline and design table for DOE 4 and 5. B-cells (1×10^3^) were seeded in 96-well plates on day 0. Flow cytometry and TRIFMA were performed on day 8. (**B**) Pareto plot of parameter effect sizes on B-cell viability in DOE 4. (**C-D**) Proliferation and viability of each sample from DOE 4 where color and mean indicated by dotted lines are specific for IL-4 pulse (**C**) and level of expression of N.40 cells (**D**). (**E**) as (**B**) but for DOE 5. (**F**) as (**C**) but for DOE 5. (**G**) as (**D**) for DOE 5. (**H**) Pareto plot of parameter effect sizes on the IgE levels measured in culture medium of DOE 4 using TRIFMA. (**I**) Measured IgG and IgE in culture DOE 4 medium, color-coded for the use of an IL-4 pulse at B-cell seeding (Day 0). (**J**) as (I) but color-coded for the use of high (N.40-high) or low (N.40-low) CD40L-expressing feeder cells. (**K**) as (H) but for DOE 5. (**L**) as (I) but for DOE 5. (**M**) as (J) but for DOE 5.

Comparing the use of purified recombinant cytokines versus feeder cells to secrete BAFF and IL-21 clearly showed identical outcomes across both viability, proliferation and Ig-secretion. For DOE 4 and 5 the highest rate of B-cell proliferation was 87- and 125-fold during the 8 days of culture, respectively. These results suggest that feeder cells delivering cytokines (except IL-4) can drive B-cell proliferation as effectively as, or even better than, purified recombinant cytokines, despite requiring additional feeder cells. For DOE 4, the highest rate was measured for the sample with 0 ng/mL BAFF and IL-4, 10 ng/mL IL-21, N.40-high, and changing just 100 µL medium compared to 200 µL. This was comparable for DOE 5, where the highest rate of B-cell proliferation was with no N.BAFF, 0 ng/mL IL-4, 500 N.IL21 cells/well, N.40-high, and changing 200 µL medium.

Notably, for the concentrations of IL-21 tested here (1-100 ng/mL), there was no significant difference for any of the readouts suggesting that under the investigated conditions using more than 1 ng/mL of IL-21 was enough to make the other components the bottleneck in terms of viability and proliferation (**Fig. S6E-F**). Considering that the previous DOE 3 with readout on day 11 suggested an IL-21 concentration of 172 ng/mL as the optimum for B-cell proliferation, there is likely a beneficial effect of using higher concentrations of IL-21 during longer cultures, however, the three samples with the highest B-cell proliferation in DOE 4 with readout on day 8 all received 10 ng/mL suggesting that 10 ng/mL is better than 100 ng/mL for proliferation in the shorter time-scale (**Fig. S6E**).

Collectively, we found that BAFF and IL-21 could effectively be supplied by feeder cells. This approach could simplify the culture system if all the necessary transgenes were incorporated into a single cell line, though it would reduce the flexibility to control their relative concentrations.

### Investigating the culture system over time measuring Ig-secretion and using flow cytometry

Next, we expanded the set-up to 6-well plates to enable flow cytometry analyses and furthermore, we harvested samples at 5 time-points during 14 days of culture (**Fig. 6A**). In addition, we expanded the measurement of Ig-secretion to include IgG1, IgG3, IgA and IgE. Here, we limited the variables to be the addition of BAFF and IL-4 and the expression level of CD40L.

**Figure 6.**
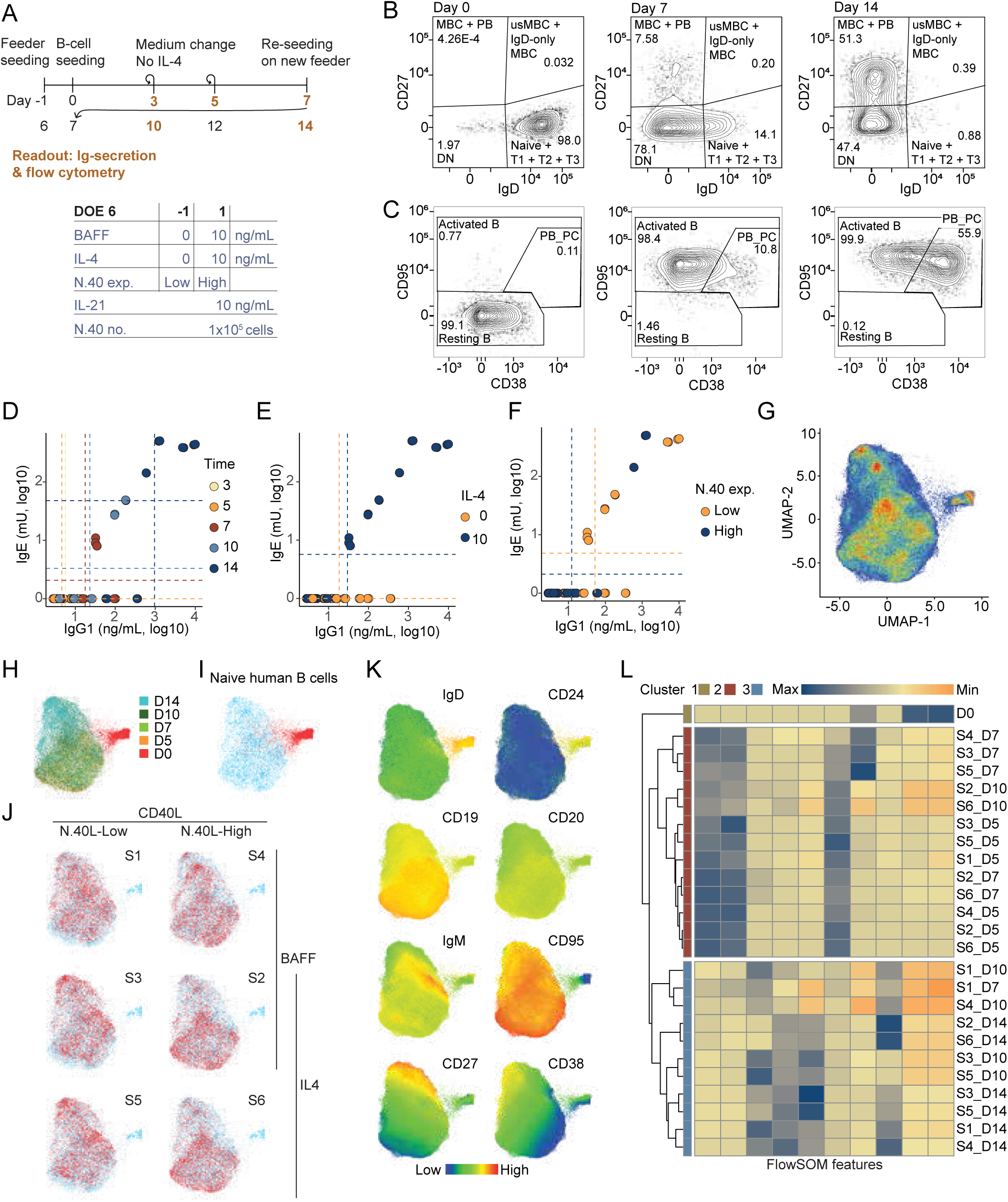
Secreted and displayed marker investigation of naïve B-cell iGB culturing. (**A**) Timeline and design table for DOE 6. B-cells (2.8×10^4^) were seeded in 6-well plates on day 0. Flow cytometry and TRIFMA were performed on day 0, 3 (TRIFMA only), 5, 7, 10, and 14. (**B-C**) Flow plotting of day 0, 7 and 14 B-cells to illustrate their activation and differentiation measured by their expression of IgD and CD27 (**B**), and CD38 and CD95 (**C**). (**D**) Measured IgG and IgE in culture DOE 6 medium by TRIFMA, color-coded as harvest day of the respective B-cells. (**E**) as (D) but color-coded for the presence of an IL-4 pulse at B-cell seeding (Day 0) (**F**) as (D) but color-coded for the use of high (N.40-high) or low (N.40-low) CD40L-expressing feeder cells. (**G**) UMAP clustering of pooled flow cytometry data. (**H**) UMAP clustering as (G) but color-coded for days in culture. (**I**) UMAP clustering as (G) color-coding the naïve B-cells seeded at day 0. (**J**) UMAP clustering as (G) but color-coded as the respective sample conditions for the DOE 6 screen. (**K**) UMAP clustering as (G) but color-coded for respective marker densities measured by flow cytometry median fluorescent intensity. (**L**) Heatmap of FlowSOM clustering analysis.

We observed the same activation and differentiation of B cells as previously seen (**Fig. 6B-C**). Modelling the secretion of Ig showed that time, IL-4, and CD40L expression influenced the secretion, whereas the addition of BAFF had no significant influence on the secretion of Ig (**Fig. 6D and S7A-D**). Furthermore, for IgG1, IgG3, and IgA, time was the most significant component, whereas IL-4 and CD40L expression were less significant (**Fig. S7E-H**). For the secretion of IgE, the addition of IL-4 with N.40-low resulted in more secretion (**Fig. 6E-F**). Notably, with time after 14 days the combination of IL-4 and N.40-high also enabled secretion of IgE at a comparable level to N.40-low. This agreed with the earlier observation where 8 days of culture in the presence of IL-4 and N.40-high did not show much IgE secretion (**Fig. 5J, M**), whereas 11 days of culture showed significant levels of IgE secretion (**Fig. 4K**).

To investigate the effect of the variables on differentiation of B cells *in vitro* in an unbiased manner, we performed UMAP clustering on the flow cytometry data (**Fig. 6G, Fig. S7I-J**). The segregation of clusters was mostly driven by time in culture, with day 0 being separate from the remaining time-points, and with day 14 being opposite to Day 5 in the big cluster along the UMAP-2 axis (**Fig. 6H-I**). Overlaying the different sample conditions showed that they grouped together in pairs of no IL-4, IL-4 & N.40-low, and IL-4 & N.40-high irrespective of BAFF (**Fig. 6J**). The panel of markers showed that the clustering of the activated samples was mostly driven by CD27, CD38, and IgM (**Fig. 6K**).

In addition, we performed FlowSOM analysis and clustered the different samples based on 10 extracted features from FlowSOM (**Fig. 6L**) (Van Gassen et al. 2015). The samples clustered into 3 large clusters, where cluster 1 was the naïve day 0 sample, cluster 2 was the early time-point samples and cluster 3 was the samples of later time-points. Interestingly, sample 2 and 6, which had received IL-4 and were cultured on N.40-high with and without BAFF, for all (except for the last time-point) clustered together with the early time-points, suggesting that IL-4 combined with N.40-high results in a slower activation and differentiation. In the opposite direction was sample 1, which had not received IL-4 and was cultured on N.40-low with BAFF, where this sample already at day 7 clustered with the late time-point samples suggesting a faster differentiation.

Taken together, our findings confirmed that time in culture is a significant determinant of the differentiation process from naïve, through activated, to memory/plasma cell phenotypes, as would be expected. However, high CD40L expression seemed to delay the differentiation process, whereas low CD40L accelerated differentiation. For Ig secretion, time was the main determinant in the default class-switching to IgG1, IgG3 and IgA, whereas class-switching to IgE required IL-4 and low-level expression of CD40L to happen within 10 days of culture. High-level expression of CD40L slowed the IgE class-switching process, and BAFF played no role in IgE class-switching.

## Discussion

Inexpensive, robust and easy-to-use primary B-cell cultures have significantly advanced our understanding of murine B-cell biology, thanks to the Nojima co-culture system and its variations (Nojima et al. 2011; Kuraoka et al. 2016). In these systems, CD40L, BAFF and IL-21 are provided by the feeder cells, whereas IL-4 is added exogenously as it needs to be provided in a pulse. Following this approach, we established similar human culture systems and employed a systematic Design of Experiments (DOE) strategy to investigate interactions among cytokines, allowing us to optimize the culture systems. Rather than isolating one parameter at a time, DOE enabled us to design and analyze multiparametric setups, uncovering key interaction effects, identifying optimal conditions, and determining the most critical parameters with minimal experiments.

Summarizing our findings, we consistently observe that IL-4 limits cell proliferation and viability, two parameters which are intimately linked. This effect can be rationalized in terms of the observed role of IL-4 in driving differentiation and class-switching to IgE. The negative effect on cell proliferation and viability can be compensated by low-level exposure to CD40L, whereas the combination of IL-4 and high CD40L is overall detrimental. CD40L in general delays differentiation, likely because it keeps the cells in an iGB-like state. In the absence of IL-4, default class-switching to IgG1, IgG3 and IgA is observed, and the main factor driving the differentiation and class-switching in this scenario is simply time.

These observations agree well with the understanding of GC responses gleaned from mouse studies, where cognate T-cell support (CD40L) signals B-cell centrocytes to return to the dark zone for continued proliferation and hypermutation, whereas a strong BCR signal is thought to govern the decision to exit the GC and differentiate into plasma cells (Lechouane et al. 2013; ElTanbouly et al. 2024). Although we have not provided any BCR stimulus in this setting, BAFF is known to be able to co-opt the BCR signaling pathway (Schweighoffer et al. 2013), but does not appear sufficient in our setup. It is also known that BAFF can support default class-switching (Castigli et al. 2005), but again it does not exert a notable effect in our cultures. IL-4, on the other hand, induces a more powerful type-specific IgE class-switching, which entails a highly risky and long-range recombination process with double-stranded breaks arresting proliferation and causing a drop in cell viability. We propose that these effects are likely amplified when CD40L levels are high, creating a conflicting combination of signals for proliferation and growth arrest, which in the presence of genomic instability, likely explains the strong negative interaction effect we observed.

The arguments to pursue better investigative tools for human primary B cells are many. Due to the ethical limitations of humans as a study-organism, mice are invaluable to obtain an *in vivo* mechanistic understanding of the complex circuitry of the immune system. However, there are obviously fundamental differences between mouse and man, in particular also for B cells. For example, in Bruton’s tyrosine kinase (BTK) deficiency, humans lack B cells entirely due to a block in the pro- to pre-B-cell transition, whereas BTK-deficient mice still produce pre- and immature B cells, highlighting divergent B-cell development (Mestas and Hughes 2004). Similar differences extend to genetic alterations, transcriptional profiles of immune-related genes like TLR7 and 9, peripheral blood leukocyte composition, and antibody isotype distributions, such as IgA1 and A2 in humans versus a single IgA in mice. Furthermore, humans possess unique Fc receptors like FcαRI and FcγRIIA/C, absent in mice, which affect humoral immune responses. These species-specific differences underscore the need for human-specific investigative tools, such as the one developed here (Mestas and Hughes 2004; Pulendran and Davis 2020; Medetgul-Ernar and Davis 2022; Gros and Casanova 2023).

To further explore human B-cell activation and differentiation, future studies could examine the role of IL-4 in facilitating germinal center (GC) B-cell exit as memory B cells, as recently proposed (Shehata et al. 2024; Duan et al. 2021). Moreover, this culture system could provide insights into processes like class-switch recombination, somatic hypermutation, and feedback loops, such as the newly described IL-12 and IFN-γ positive feedback loop in T-independent activation, which may also apply to T-dependent activation (Elsner, Smita, and Shlomchik 2024). In addition, the emerging evidence of synergistic effects between BCR and CD40L stimuli (Nova-Lamperti et al. 2017; Rauert-Wunderlich et al. 2018) and between IL-21 and CD40L (Ding et al. 2013) could be investigated further. Our findings suggest similar interactions between IL-4 and CD40L, positioning CD40L signaling as a central interactor. While CD40L-CD40 signaling is best known for its NF-κB activation via canonical and non-canonical pathways, it can also engage MAPK, PI3K, and PLCγ signaling pathways, offering multiple regulatory nodes (Elgueta et al. 2009). In our experiments, IL-21 was included in all cultures due to its established importance in prior studies (Avery et al. 2008; Kim et al. 2024). However, future work could further investigate its specific role by comparing cultures with and without IL-21. Finally, our system may serve as a foundation for producing patient-derived therapeutic antibodies, offering a platform for antibody development.

In conclusion, our study highlights the power of human primary B-cell culture systems, systematically characterized and optimized using a DOE approach, to uncover key cytokine interactions, such as those between IL-4 and CD40L, and their roles in B-cell activation and differentiation. This user-friendly platform provides a robust foundation for advancing our understanding of human immune mechanisms and supports future applications in immunological research and therapeutic development.

## Materials and Methods

### Cell lines

HEK293T (Lenti-X 293T) cells (Takara Bio, cat. #632180) were cultured in DMEM supplemented with 5% fetal bovine serum (FBS) and 1% penicillin/streptomycin (P/S). MS-5 cells (DSMZ, cat. #ACC 441) and the primary Normal Human Dermal Fibroblasts (NHDF) cells NHDF03 (Promocell, Lot #4090701.2) and NHDF15 (CellSystems, Lot #03410) were cultured in RPMI 1640 supplemented with 10% FBS and 1% P/S. Cells were cultured at 37°C with 5% CO_2_.

### Plasmids

pCCL-PGK-hTert-IRES-Hygro and pCCL-PGK-IL4-IRES-Puro were generated by linearization of pCCL-PGK-eGFP (Jakobsen et al. 2009) using BamHI and XhoI (Thermo Fisher), following insertion of either two fragments amplified from pLV-hTERT-IRES-hygro (Addgene #85140, kindly provided by Tobias Meyer) or one fragment amplified from pAIP-hIL4-co (Addgene #74169, kindly provided by Jeremy Luban) using primers listed in Supplementary Table 1. pCCL-PGK-BAFF-IRES-Puro was made by inserting a fragment amplified from tnfsf13b-in-pdonr201 obtained from Harvard Plasmid Bank into first linearized pLV_PGK-mCherry-BHGpA_CMV-eGFP-WHV and then subsequently into BamHI-digested pCCL-PGK-MCS-IRES-Puro (Gao et al. 2022) using primers listed in Supplementary Table 1. pCCL-PGK-CD40LG-IRES-Puro was constructed as described in (Nielsen et al. 2024). pCCL-PGK-IL21-IRES-Puro was generated by linearization of pCCL-PGK-MCS-IRES-Puro using BamHI and insertion of a PCR amplified fragment (synthesized by Twist Biosciences) using primers listed in Supplementary Table 1. Assembly of all constructs was performed using NEBuilder HiFi DNA Assembly Master Mix (New England Biolabs) according to manufacturer’s instructions.

### Lentiviral vector production

For lentiviral vector production, 4 × 10^6^ HEK293T cells were seeded and transfected 24 hours later with 3 μg pRSV-REV, 13 μg pMDIg/pRRE, 3.75 μg pMD.2G and 13 μg of the desired lentiviral transgene vector. Transfection was performed using calcium phosphate buffers. Medium was changed 24 hours after transfection, and 48 hours after transfection, viral supernatants were harvested and filtered through a sterile filter (0.45 μm).

### Generation of feeder cells

Cells were transduced by seeding 1 × 10^5^ cells in a 6-well plate in RPMI medium with 10% FBS, 1% P/S and 8 µg/mL polybrene and then adding the lentiviral vector preparation to a total volume of 2 mL. The NHDF cells were immortalized by transduction of lentiviral vectors carrying vector RNA encoded from pCCL-PGK-hTert-IRES-Hygro inducing expression of human telomerase reverse transcriptase (*hTert*) and hygromycin B phosphotransferase (*Hygro*). The cells underwent 1 week of hygromycin selection. To engineer expression of BAFF, CD40L, IL-4 and IL-21, the NHDF and MS-5 cells were transduced with the respective lentiviral vectors encoding the gene coupled through an IRES sequence to the puromycin resistance gene; pCCL-PGK-GOI-IRES-Puro. The cells were transduced in triplicates with MOIs of ∼1, ∼3 and ∼5 to give a larger range of expression levels. The MOI was estimated from same batch pCCL-PGK-eGFP lentiviral vector preparations in transducing the respective cell line using a Fluorescence Titering Assay. The cells underwent 1 week of puromycin selection. Next, to expand single cells, limiting dilution was used to seed ∼7.5, ∼15 or ∼30 cells per well in 96-wells in 100 µL RPMI with 30% FBS and 1% P/S. Half the medium was replenished once a week for 6-10 weeks until the cells could be transferred to a 24-well and expanded as normal. Notably, each single well was observed for growth of only a single colony before qualifying for expansion.

### Purification of primary human B cells

To purify human naïve B cells, PBMCs were first isolated from human buffy coats, obtained from anonymized healthy donors at the blood bank of Aarhus University Hospital, Denmark. Human buffy coat was first mixed 1:1 with PBS, PBMCs were then isolated by a Ficoll-Paque PLUS (GE Healthcare) density gradient centrifugation using SepMate tubes (Stemcell Technologies) centrifuged at 1200 g for 10 min at 20 °C. The PBMCs were subsequently washed in PBS, centrifuged at 500 g for 10 min at 20 °C, then incubated in PBS with 27 mM EDTA for 5 min at RT before centrifugation at 300 g for 8 min at 20 °C followed by another wash in PBS. The clean PBMCs were resuspended in 5 mL MACS buffer (PBS, 2% heat-inactivated Fetal Bovine Serum (FBS), 2 mM EDTA) at RT, before being counted and 400 ×10^6^ cells taken aside for the MACS purification protocol. For the magnetic isolation of human naïve B cells, the PBMCs were first incubated 5 min on ice with Human BD Fc block (Clone Fc1, BD, 564220) before being treated with the Miltenyi Biotech Human Naïve B Cell Isolation biotin-antibody cocktail (130-091-150) for 30 min. The samples were then diluted with MACS buffer and centrifuged at 200 g 10 min at 4°C. The pellet was resuspended in 1 mL MACS and incubated 20 min with Miltenyi Biotec anti-biotin magnetic beads, pre-diluted in 2 mL MACS buffer. The cell suspension was loaded onto pre-wetted Miltenyi Biotec LS columns through 70 μm cell strainers and washed 2 times with 3 mL MACS buffer. The identity of the collected flowthrough of untouched human naïve B cells (CD19^+^, CD20^+^, CD22^+^, CD27^−^, IgD^+^) was validated using flow cytometry. Cell aliquots of human B cells were frozen in freeze media (90% FBS, 10% DMSO) using the Corning CoolCell containers as recommended by the manufacturer.

Human CD19^+^ B cells were purified from buffy coats using the StraightFrom Buffy Coat CD19 MicroBead purification Kit (Miltenyi Biotec, cat. #130-114-974) following the manufactures instructions. Purification was subsequently assessed by flow cytometry using a panel of B-cell markers (CD19^+^, CD20^+^, CD24^+^, CD27^+^, CD38^+^ and IgD^+^). Cells were frozen in freeze media (90% FBS, 10% DMSO).

### Inducible Germinal Center B cell (iGB) co-culture system

Feeder cells, as population or clone, were cultured at least 8 days before being seeded and incubated in B-cell medium (BCM; RPMI-1640 [+] L-Glutamine supplemented with 55 µM 2-Mercaptoethanol, 1% P/S, 10 mM HEPES, 1 mM Sodium Pyruvate (all Invitrogen) and 10% FBS. The following day (day 0), human pan or naïve B cells were seeded onto the feeder cells in BCM supplemented with IL-4, IL-21 and BAFF (Peprotech, 200-04, 200-21 and 310-13, respectively) in a final culture concentration as dictated by the experiment. On day 3 and 5, half the culture volume of media was exchanged with fresh BCM similarly supplemented with IL-21 and BAFF. On day 7, B cells were harvested then subjected to the analysis of interest or re-seeded on fresh feeder cells. Culture supernatants were stored with 0.1% Sodium Azide for later TRIFMA analysis of secreted antibody isotypes as described below.

The above was performed for all the culture experiments with DOE 1 through 5 experiments using pan (CD19^+^) B cells and DOE 6 using naïve B cells.

### Flow cytometry

For viability and proliferation assays, the samples were mixed after which 97.5 μL cell suspension was transferred to a V-bottom 96-well plate with 2.5 μL 50 μg/mL propidium iodide. The number of live and dead cells in 30 μL was counted using a NovoCyte 2100YB Flow Cytometer (Agilent Technologies) as done in (Rovsing et al. 2023).

Cell-suspensions of interest were prepared in flow cytometry (FC) buffer (PBS with 2% FBS and 1 mM EDTA), either being freshly MACS-purified human B cells to validate purity or B cells harvested from human iGB cultures for identity analysis. Human PBMCs were thawed for the respective compensation controls. One hundred µL of sample was initially pre-incubated 5-10 min with 20 µL Fc Block dilution. This was followed by staining with 100 µL fluorophore-conjugated antibody panel of choice (see **Supplementary Table 2**) diluted in 1:1 FC buffer with Brilliant Stain buffer (DB, 563794). After 30 min of staining on ice, the cells were washed and subsequently resuspended in the FC buffer. Before analysis, all samples were strained through a 100 µm filter to avoid aggregates. The stained cell solutions were analyzed on NovoCyte Quanteon 4025 (Agilent) equipped with 4 laser (405, 488, 561 and 640 nm) and 25 detectors. The acquired data were analyzed using FlowJo v. 10.10.0 (BD).

### Flow Cytometry Clustering analysis

Live, CD19^+^ singlets from every sample were down sampled to 10,000 events using the DownSampleV3.3.1 plugin for FlowJo v10.10.0. These were concatenated and clustered into 10 clusters using Self-Organizing Maps with the FlowSOM v4.1.0 FlowJo plugin (Van Gassen et al. 2015). An independent UMAP clustering was made using the EmbedSOM v2.2.0 FlowJo plugin (Kratochvíl, Koladiya, and Vondrášek 2019).

### ELISA and Time-Resolved Immuno-Fluorometric Assay (TRIFMA)

To verify and measure the cytokine production from feeder cells, the cells were seeded in 6-well plates at the indicated densities in 2 mL RPMI medium. After 24 hours, medium was changed to 3 mL BCM medium and after 96 hours culture supernatants were harvested. ELISA was performed on dilutions of culture supernatant in duplicates using the following kits: Human BAFF/BLyS/TNFSF13B DuoSet ELISA (#DY124-05), Human IL-4 DuoSet ELISA (#DY204-05) and Human IL-21 DuoSet ELISA (#DY8879-05) (all from R&D Systems) according to manufacturer’s instructions.

To quantify secretion of antibody isotypes, TRIFMA was performed on culture supernatants. FluoroNunc MaxiSorp 96-well plates were coated with 100 µL per well of target-binding antibody diluted in PBS. 0.3 µg/mL Donkey anti-human IgG (Jackson ImmunoResearch, 709-005-098), 2.5 µg/mL Mouse Anti-human IgG1 (SouthernBiotech, 9052-01), 1 µg/mL Mouse Anti-human IgG3 (SouthernBiotech, 9210-01), 1 µg/mL Goat Anti-Human IgA (SouthernBiotech, 2053-01), or 2.5 µg/mL anti-human IgE (SouthernBiotech, 9240-01), all incubated o/n at 4 ℃. Wells were emptied and blocked with 200 µL TBS (Tris-buffered saline (137 mM NaCl, 2.7 mM KCl, 25 mM Tris/Tris-HCl) containing 0.09% (w/v) sodium azide (Fisher Scientific, BP2471-500)) with 1% BSA (Sigma A 4503) for 1 hour at RT and washed 3 times using TBS with 0.05% (v/v) Tween-20 (TBS/TW). All samples were diluted 1/5 in TBS/TW/0.1% BSA and added in duplicates to wells for 2 hours incubation at RT (1 h for pan IgG). Standard curves were made with TBS/TW/0.1% BSA diluting either; human IgG (gammanorm, 478393) to 2000 ng/mL and further 7 5-fold dilutions; human IgG1 (SouthernBiotech, 0151-K) to 100 ng/mL and further 10 2-fold dilutions; human IgG3 (SouthernBiotech, 0153-L) to 100 ng/mL and further 10 2-fold dilutions; human IgA (SouthernBiotech, 0155-L) to 500 ng/mL and further 10 2-fold dilutions; or for IgE, human plasma diluted 3-fold and further 10 times 2-fold. Following the incubation, wells were washed three times with TBS/TW before exposure at RT with appropriate biotinylated detection antibody; Goat anti-human IgG (H+L) (SouthernBiotech, 2016-08) diluted in TBS/TW/0.1% BSA to 0.1 µg/mL, incubated 1 hour for pan IgG detection, likewise for IgG1- and IgG3-detection, but with the latter exposed for 2 hours, both diluted in TBS/TW; Goat anti-human IgA (Invitrogen, A18791) diluted in TBS/TW to 0.5 µg/mL incubated 2 hours; Mouse anti-human IgE (SouthernBiotech, 9250-08) diluted in TBS/TW to 0.2 µg/mL exposed for 2 hours. Wells were washed three times in TBS/TW and incubated 30 min at RT with Eu^3+^-labeled streptavidin (PerkinElmer, 1244-360), diluted to 1 µg/mL in TBS/TW with 25 µM EDTA. Lastly, wells were washed three times with TBS/TW and added 200 µL Enhancement solution (Ampliqon). Plates were shaken 5 min and measured on a time-resolved fluorometry plate reader (Victor X5 Perkin Elmer).

### Design of Experiments (DOE)

The experimental setup for each DOE was manually designed based on Full Factorial and Box Behnken designs. Additional replicates of specific conditions were included in each design to enable more precise estimates of the model fit R^2^. The experimental run order was randomized to avoid bias. Backward Elimination was used such that only variables with a statistically significant (p < 0.05) coefficient were included in the linear model. Across all linear models the minimum adjusted R^2^ was 0.67 and the average was 0.85. The DOEs were analyzed in R v4.2.2 using the base *lm* function, the pid package v0.50 and custom code. Data, linear models and scripts for all DOE experiments can be found on Github (https://github.com/arovsing/DOE_iGB-culture). For further information, the coursera course ‘Experiment for Improvement’ created by Kevin Dunn contains all information needed to design and run DOEs.

## Abbreviations

BCM: B-Cell Medium
BCR: B-Cell Receptor
CD40L: CD40-ligand
CSR: Class-Switch Recombination
DN: double-negative
DOE: Design of Experiments
GC: Germinal Center
IFN: Interferon
Ig: Immunoglobulin
iGB: inducible Germinal Center B cell
IL: Interleukin
MBC: memory B cell
NHDF: Normal Human Dermal Fibroblasts
PB: plasma blast
PC: plasma cell
SHM: Somatic Hyper Mutation
usMBC: unswitched MBC.

## Acknowledgements

This work was funded by grants from the Independent Research Fund Denmark to JGM (9039-00173B) and SED (Sapere Aude Research Leader grant; 9060-00038). Additional funding was provided by Aase og Ejnar Danielsens Fond and the NEYE foundation. We are grateful to the FACS Core at AU for expert assistance with flow cytometry experiments.

## Conflict of interest statement

JGM is a member of the Scientific Advisory Board of Nvelop Therapeutics. The company was not involved in the present study. The remaining authors declare no commercial or financial conflict of interest.

## Ethics approval statement

B cells were purified from buffy coats, obtained through the blood bank at Aarhus University Hospital, from anonymized healthy volunteers, in accordance with the Danish Health Care Act (Sundhedsloven).

## Author contributions

Conceptualization, ABR, KG, JGM and SED; Methodology, ABR, KG, JGM and SED; Formal analysis, ABR; Investigation, ABR, KG, IHN and LJ; Writing – Original Draft, ABR, KG and SED; Writing – Review & Editing, ABR, KG, IHN, LJ, JGM and SED; Visualization, ABR and KG; Supervision, JGM and SED; Project Administration, ABR and KG; Funding Acquisition, JGM and SED.

## Supplementary figure legends

**Figure S1.**
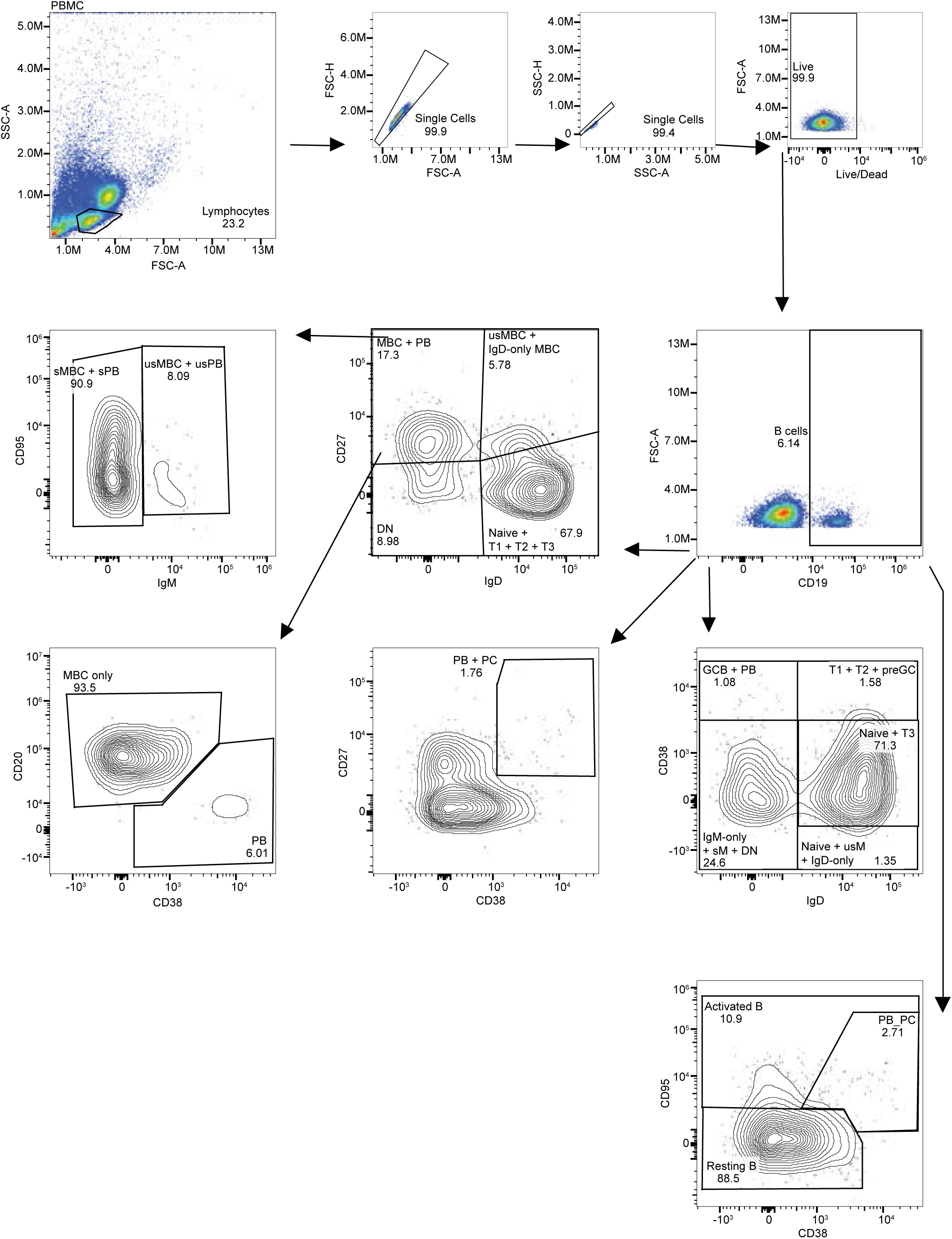
Flow cytometry gating strategy. Stepwise gating of human PBMCs, exemplifying the lymphocyte/singlet-FSC/singlet-SSC/live/CD19^+^ hierarchy used in subsequent analyses.

**Figure S2.**
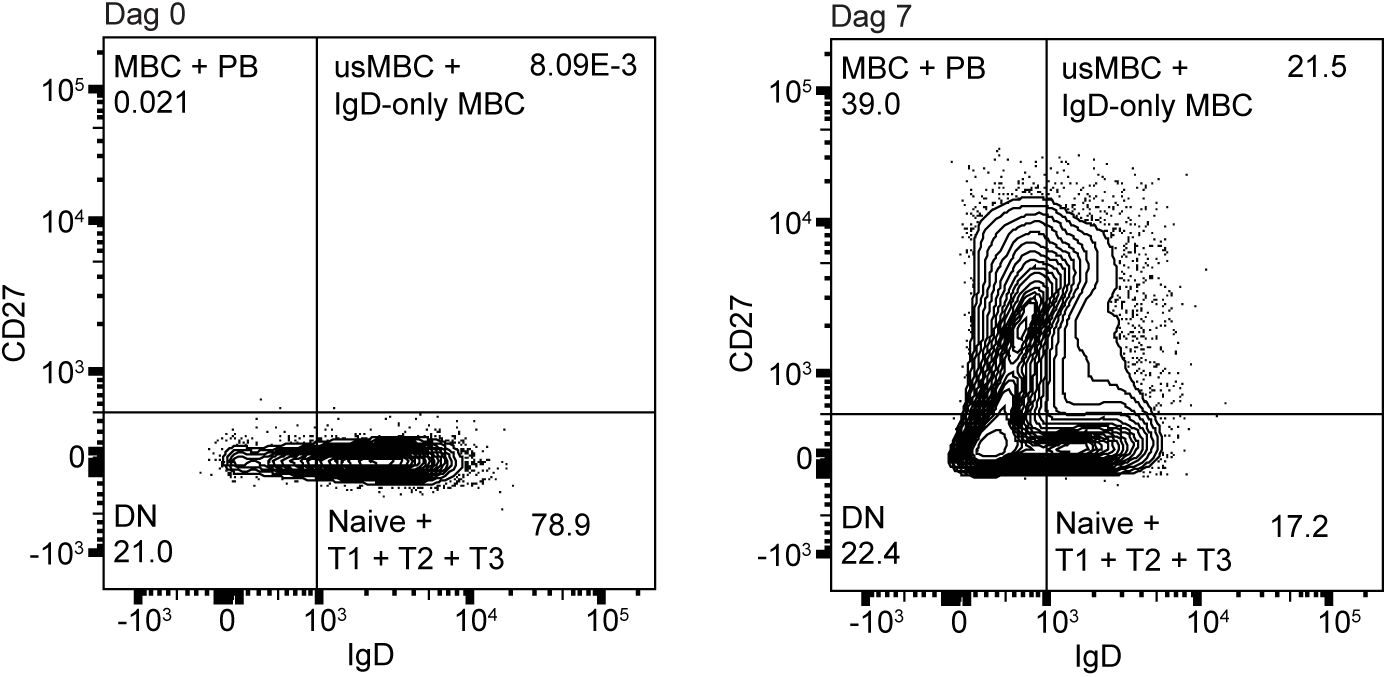
Naïve B-cell activation with NHDF03 feeder cells. Flow cytometric comparison of naïve B cells at Day 0 vs. Day 7 when co-cultured with NHDF03-hTert-CD40L cells, illustrating progressive upregulation of B-cell activation markers.

**Figure S3.**
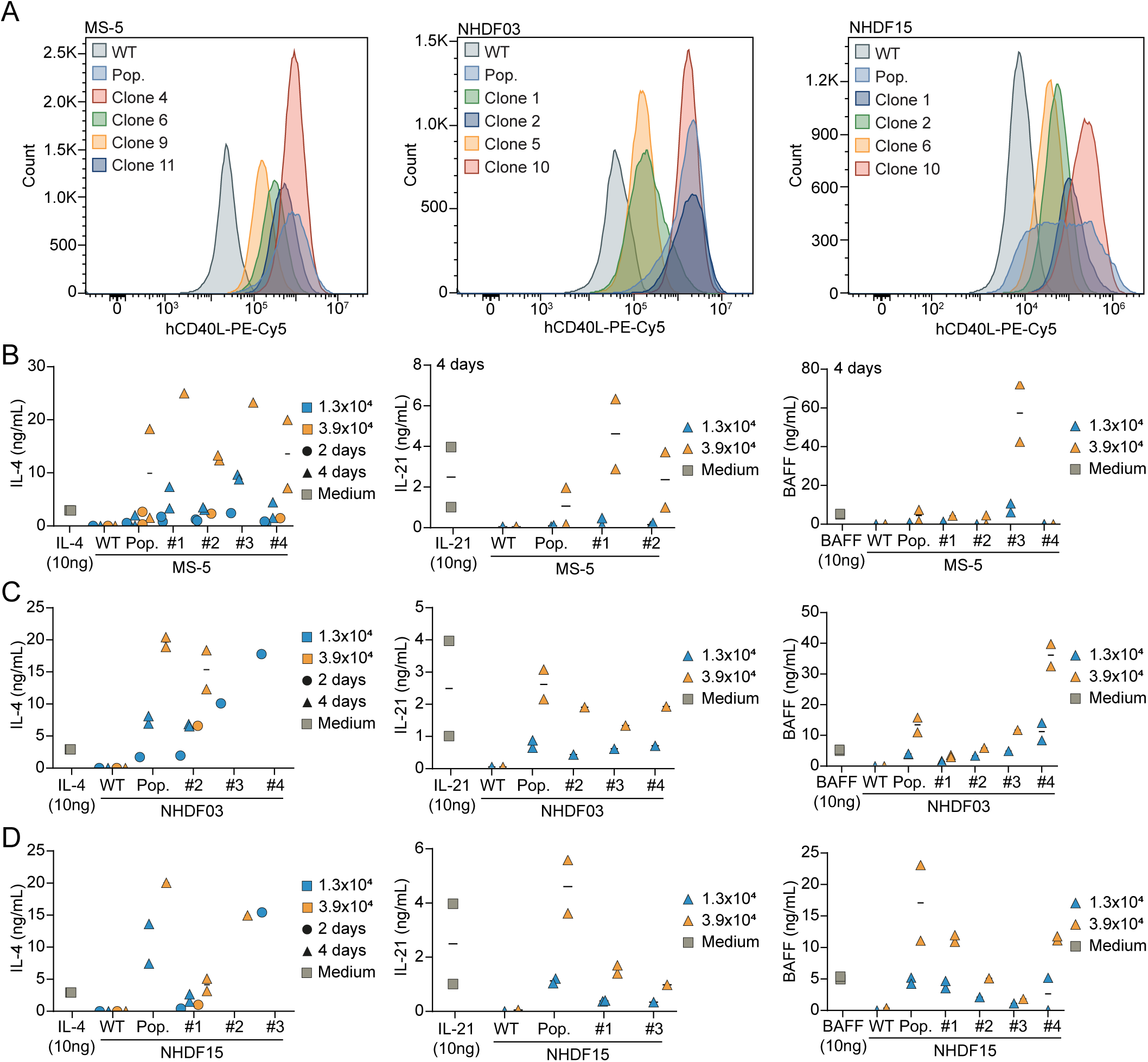
Generation of feeder cells from murine MS-5 and human NHDF03/NHDF15 lines. (**A**) CD40L display of MS-5, NHDF03 and NHDF15 measured by flow cytometry. (**B-D**) ELISA confirmation of IL-4, IL-21, and BAFF secretion by transgenic MS-5 (B), NHDF03 (C), and NHDF15 (D) feeder cells.

**Figure S4.**
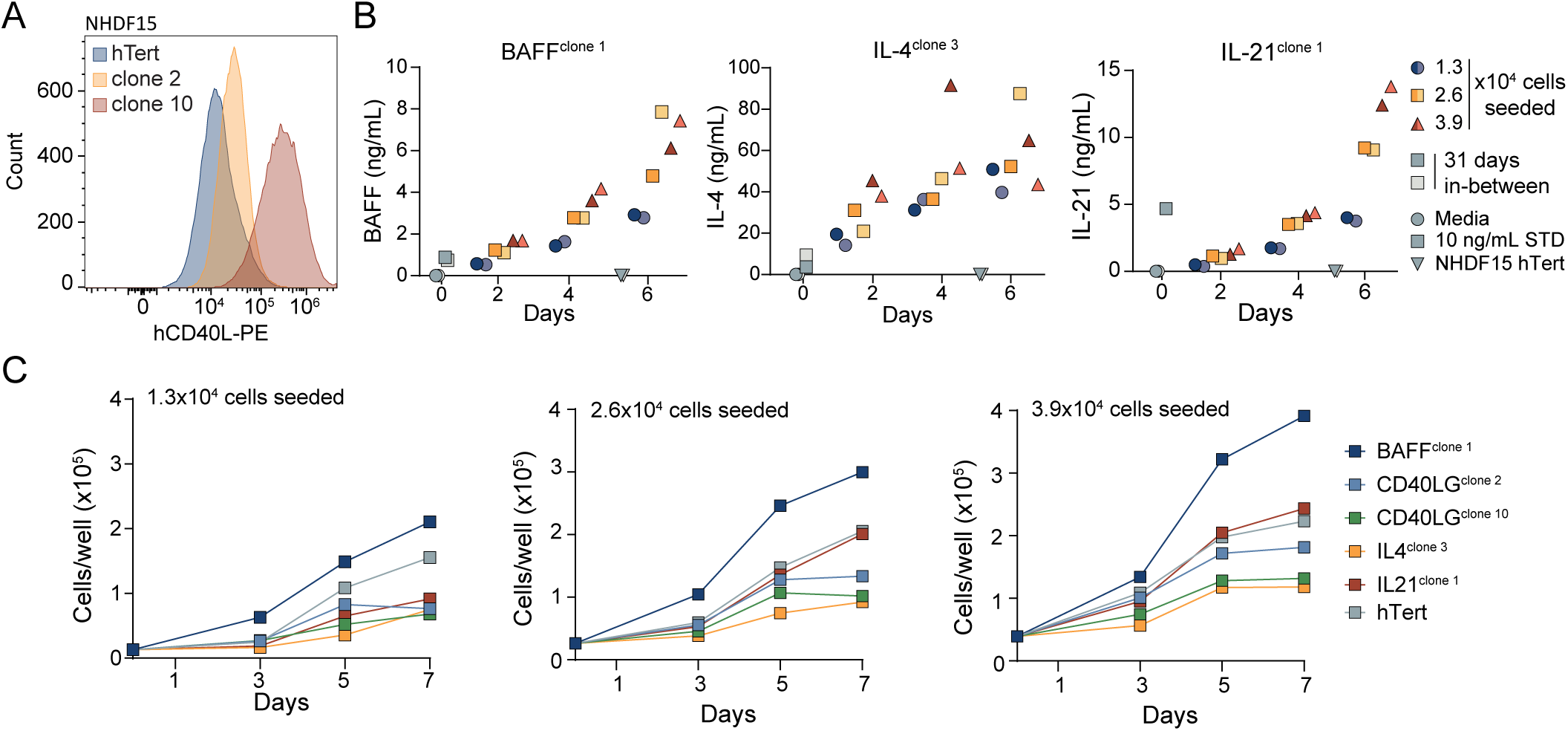
Characterization of NHDF15 transgenic clones. (**A**) CD40L expression on NHDF15 clones 2 (N.40-low) and 10 (N.40-high), as well as hTert negative controls, by flow cytometry. (**B**) ELISA-based measurement of IL-4, IL-21, and BAFF secretion over time. (**C**) Longitudinal assessment of feeder-cell proliferation by flow cytometry.

**Figure S5.**
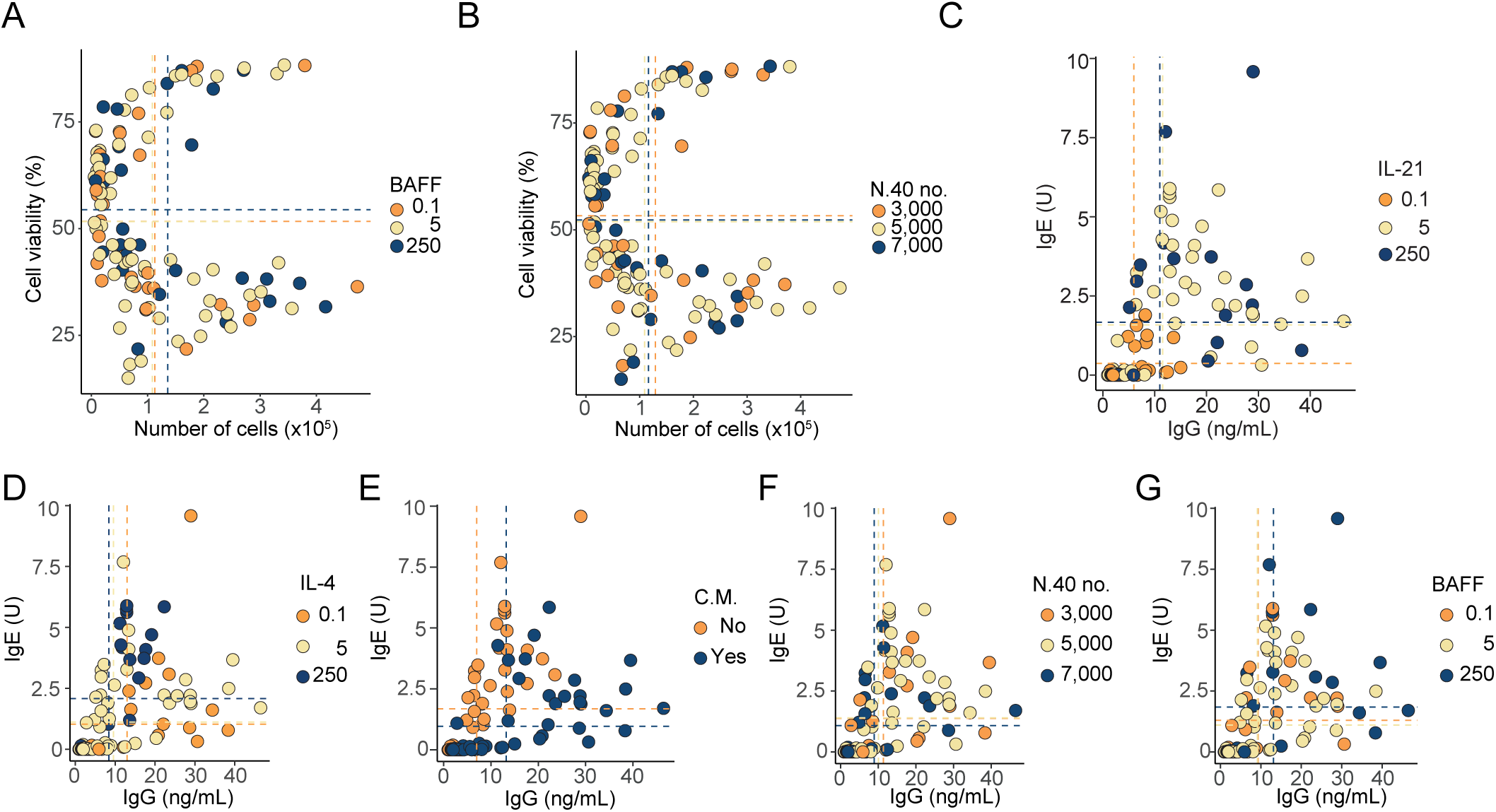
DOE 3 supplementary findings. (**A-C**) Proliferation and viability of each sample from DOE 3 where color and mean indicated by dotted lines are specific for BAFF (**A**), number of N.40 cells (**B**), and IL-21 (**C**). (**D-G**) Measured IgG and IgE in culture medium, color-coded for the level of IL-4 (**D**), change of medium (**E**), number of N.40 cells (**F**), and level of BAFF (**G**). C.M. indicates medium change.

**Figure S6.**
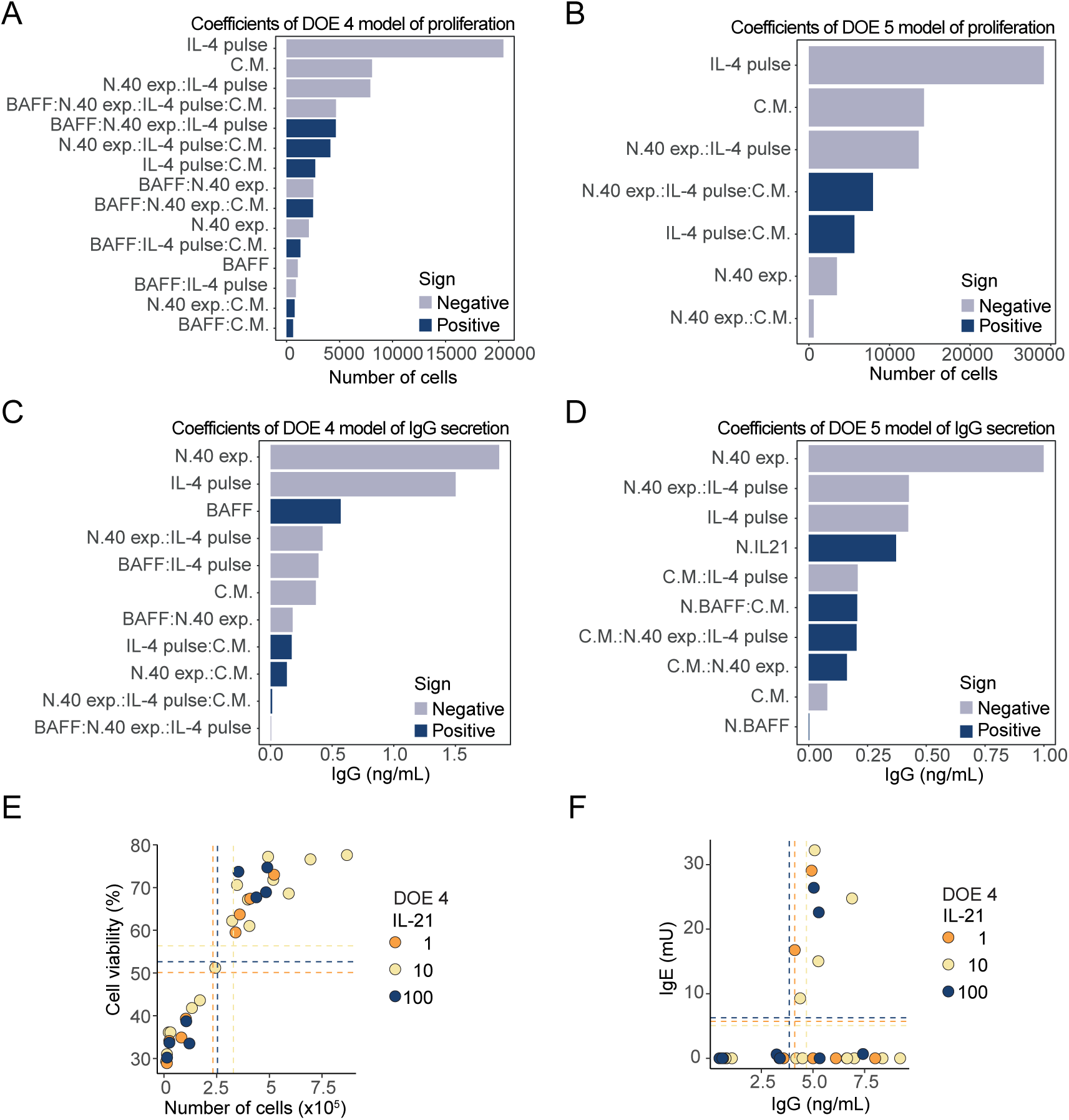
Supplementary finding for DOE 4 and 5 comparison of cytokine delivery method. (**A-B**) Pareto plot of parameter effect sizes on B-cell proliferation in DOE 4 (**A**) and DOE 5 (**B**). (**C-D**) Pareto plot of parameter effect sizes on IgG detection in the medium in DOE 4 (**C**) and 5 (**D**), measured by TRIFMA. (**E**) Raw data plotting of B-cell number and viability after 8 days of iGB culturing, with dotted lines showing the mean of the respective factor. Here color-coding for the number of seeded IL-21-expressing feeder cells in DOE 4. (**F**) As (**E**) but for measured IgG and IgE.

**Figure S7.**
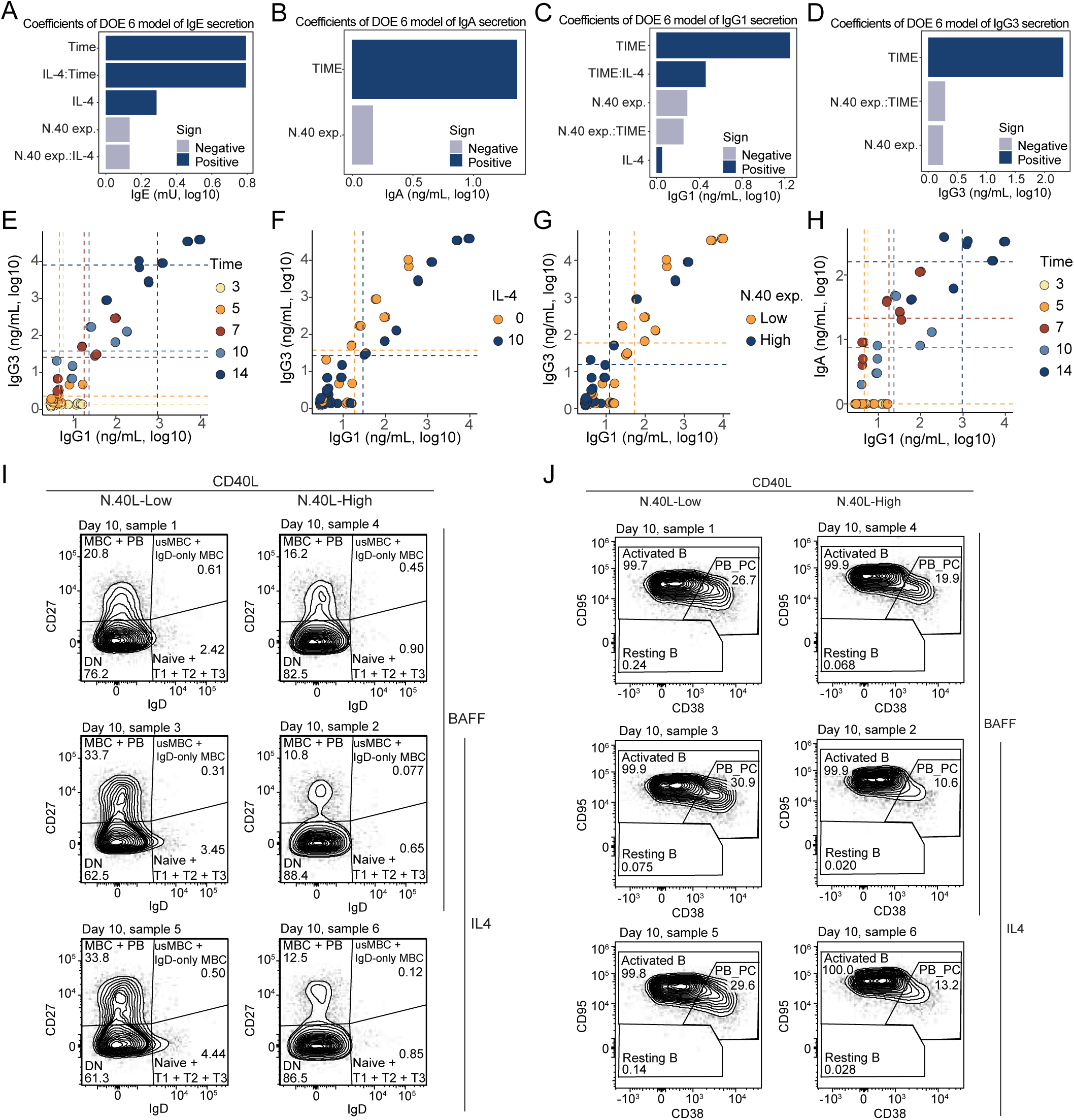
DOE 6 supplementary findings. (**A-D**) Pareto plot of parameter effect sizes on IgE (**A**), IgA (**B**), IgG1 (**C**), and IgG3 (**D**) detection in the medium of DOE 6 cultures measured by TRIFMA. (**E-G**) Measured IgG1 and IgG3 in culture medium of each sample from DOE 6 where color and mean indicated by dotted lines are specific for time (**E**), level of IL-4 (**F**), N.40 expression (**G**). (**H**) As (**E**) but for measuring IgG1 and IgA. (**I**) Flow plotting IgD vs. CD27 of CD19^+^ live singlet lymphocytes after 10 days of naïve B-cell iGB culturing to illustrate the difference of their activation and differentiation depending on the stimuli provided, with the feeder-cell expression level of CD40L being a very apparent contributor. (**J**) as (**I**) but plotting CD38 vs. CD95.

