## Supplemental Tables for "Multiparametric optimization of human primary B-cell cultures using Design of Experiments"

### **Supplementary Table 1 – Primers used for Gibson assembly**

| **Primer** | **Sequence (5’ -> 3’)** |
| --- | --- |
| pCCL-PGK-hTert-IRES-Hygro | |
| hTERT fragment, forward | GGCCTTTCGACCTCTAGCGGTGTCGTGAGGATCCATGC |
| hTERT fragment, reverse | AGGGAAACCGTTGCTAGC |
| Hygro fragment, forward | AAGCTAGCAACGGTTTCCCT |
| Hygro fragment, reverse | CCAGAGGTTGATTATCGGAATTCCCGCCCGGGCTATTCCTTTGCCCTC |
| pCCL-PGK-IL4-IRES-Puro | |
| IL4-IRES-Puro fragment, forward | CTCCGGGCCTTTCGACCTCTAGCGGTCTAGAAGCGCTGGATCC |
| IL4-IRES-Puro fragment, reverse | CCAGAGGTTGATTATCGGAATTCCCTCTCGAGATTAATCAGGCACCGGGC |
| pLV-PGK-mCherry-BHGpA_CMV-TNFSF13B-WHV | |
| BAFF fragment 1^st^, forward | AGTGAACCGTCAGATCTTTTCTAGACACCATGGATGACTCCAC |
| BAFF fragment 1^st^, reverse | GGTTGATTATCGGAATTCCCTCGAGTCACAGCAGTTTCAATGC |
| pCCL-PGK-BAFF-IRES-Puro | |
| BAFF fragment 2^nd^, forward | TGTCGACGATATCTTCGAAGGACACCATGGATGACTCC |
| BAFF fragment 2^nd^, reverse | ACGGCCGCTATGCTTTACTGGGATCCTCACAGCAGTTTCAATGC |
| pCCL-PGK-IL21-IRES-Puro | |
| IL21 fragment, forward | TGTCGACGATATCTTCGAAGACGCGTGCCACCATGAGATC |
| IL21 fragment, reverse | ACGGCCGCTATGCTTTACTGTTAATTAACTAAGAGTCCTCTGACCCATG |

### **Supplementary Table 2 - Flow Cytometry Antibodies**

| **Anti-human antibody** | **Dilution** | **Clone, Company, Cat#** |
| --- | --- | --- |
| Mouse α-CD19-PE | 1:250 | Clone HIB19, BD, 555413 |
| Mouse α-CD20-BV421 | 1:250 | Clone 2H7, BD, 562873 |
| Mouse α-IgD-PE/Cy7 | 1:250 | Clone IA6-2, BD, 561314 |
| Mouse α-CD8-PerCP/Cy5.5 | 1:250 | Clone SK1, BD, 565310 |
| Mouse α-CD4-AF647 | 1:250 | Clone RPA-T4, BD, 557707 |
| Mouse α-CD14-BV605 | 1:250 | Clone M5E2, BD, 564055 |
| Mouse α-CD16-FITC | 1:250 | Clone 3G8, BD, 555406 |
| Human BD Fc Block | 1:50 | Clone Fc1, BD, 564220 |
| Viability Dye eFluor 780 | 1:2000 | eBioscience, 65-0865-14 |
| Mouse α-CD19-BV421 | 1:250 | Clone HIB19, BD, 562440 |
| Mouse α-IgG-PE/Cy5 | 1:50 | Clone G18-145, BD, 551497 |
| Mouse α-IgM-BV605 | 1:200 | Clone G20-127, BD, 562977 |
| Goat α-IgA-FITC | 1:400 | Polyclonal, Invitrogen, H14101 |
| Mouse α-CD20-AF700 | 1:50 | Clone 2H7, BD, 560631 |
| Mouse α-CD24-PE/CF594 | 1:50 | Clone ML5, BD, 562405 |
| Mouse α-CD27-BV786 | 1:250 | Clone L128, BD, 563327 |
| Mouse α-CD38-APC | 1:15 | Clone HIT2, BD, 555462 |
| Propidium Iodide | 1.25 µg/mL | Invitrogen, BMS500PI |
| Mouse α -CD154 (CD40L)-PE/Cy5 | 1:50 | Clone 24-31, Invitrogen, 15-1548-42 |
